# Sodium Ions Regulate GPCR Activation by Remodeling Allosteric Coupling Networks and Hydration Patterns

**DOI:** 10.64898/2026.03.27.714850

**Authors:** Lisa Schmidt, Bert L. de Groot

## Abstract

Sodium ions (*Na*^+^) are key modulators of G-protein coupled receptor (GPCR) function, yet their mechanistic role remains incompletely understood. Here, we reveal a novel mode of *Na*^+^-mediated inactivation in the dopamine D2 receptor (DRD2), where *Na*^+^ reshapes long-range allosteric coupling networks and disrupts a continuous internal water column essential for activation. Using extensive molecular dynamics simulations and alchemical free energy calculations, we show that *Na*^+^ induces inactive-like residue interactions in the active state and triggers the formation of a distinct hydration gap. We also identify previously unreported *Na*^+^ binding sites and quantify their impact on the active-inactive state equilibrium by thermodynamic scanning. These findings provide mechanistic insights into *Na*^+^-driven allosteric regulation of GPCRs and highlight new opportunities for drug design targeting ion-sensitive receptor states.

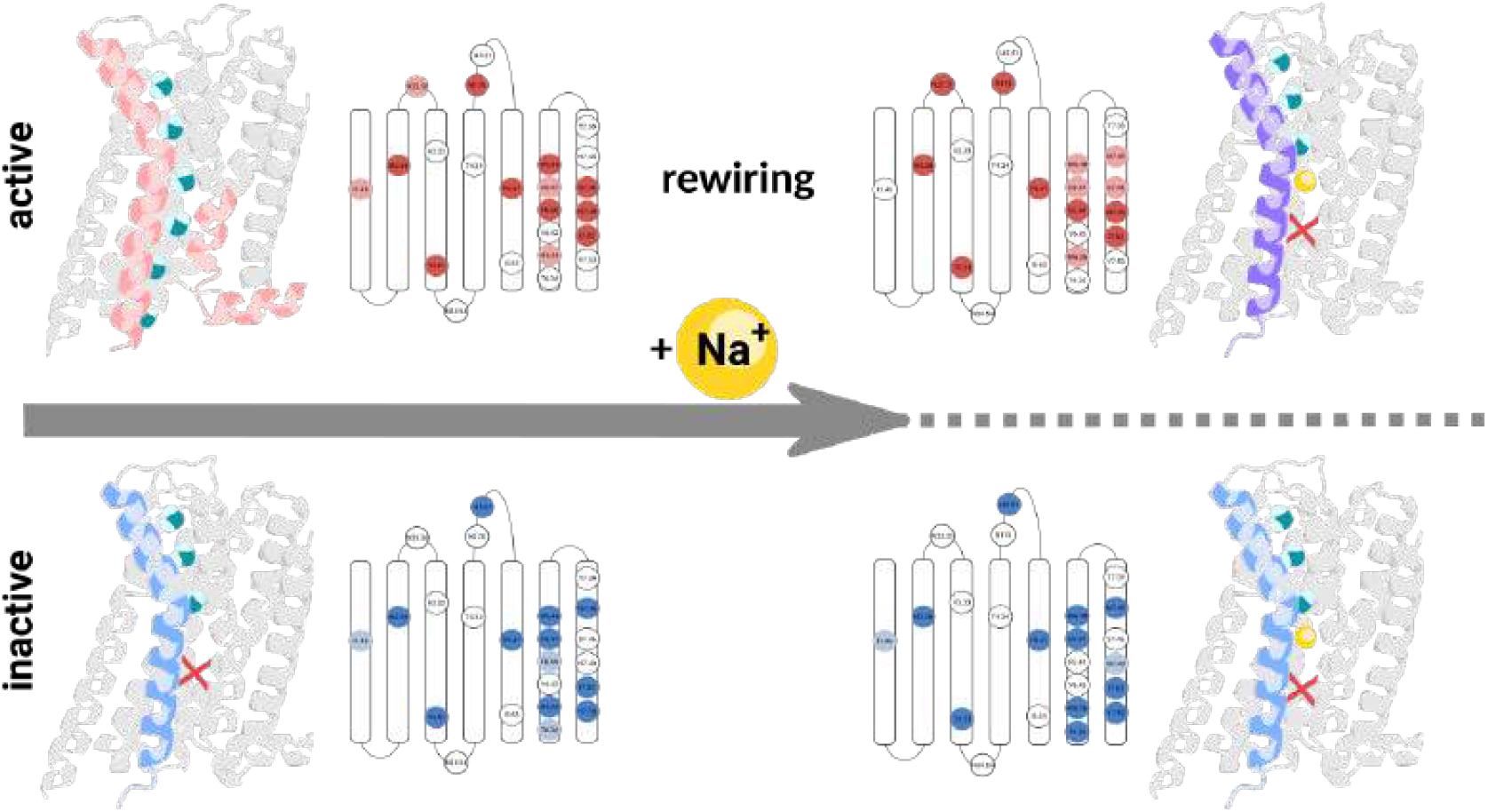

## 1 Introduction

G-protein coupled receptors (GPCRs) act as “biological computers” and are able to integrate various external signals and to switch between active (on) and inactive (off) states. ^1,2^ They respond to various effectors, including ligands, ions, lipids, and water.^1–14^ A key feature of GPCR regulation is their allosteric nature: effector binding at one site can influence distant receptor regions, thereby enabling dynamic responses to complex signals.^1,2,12,14–18^

While the separate effects of allosteric modulators like sodium ions, ligands and mutations on GPCRs are well-studied, ^1–8,13,14,19^ their collective role in modulating receptor function remains unclear. In particular, although sodium ions (*Na*^+^) are known to stabilize the inactive state of class A GPCRs like DRD2,^3–8^ the full extent of their mechanistic role in modulating receptor dynamics and signaling remains elusive.

In this study, we demonstrate a novel *Na*^+^-dependent mechanism of GPCR regulation. Using DRD2 as a model, we show that *Na*^+^ not only reinforces inactive states but actively rewires allosteric coupling networks and interrupts critical hydration channels necessary for receptor activation. Our approach integrates long-timescale simulations with free energy calculations^19–22^ and network-based coupling analysis to capture dynamic, system-wide changes in receptor behavior.

## 2 *Na*^+^ interactions differ in the active and inactive state

The interaction of sodium ions (*Na*^+^) with the DRD2 receptor affects ligand binding and stabilizes the inactive state. ^5,6,13,14,23–26^ In our simulations, *Na*^+^ interacts with different transmembrane helices (TM2, TM3, TM6, and TM7),^5–7,23–27^ with key interactions occurring at components of the allosteric *Na*^+^ binding pocket^5,6,23–26^ such as D2.50_80_ and S3.39_121_ (fig. 1b and SI section *Na*^+^ contact sites).

**Figure 1:**
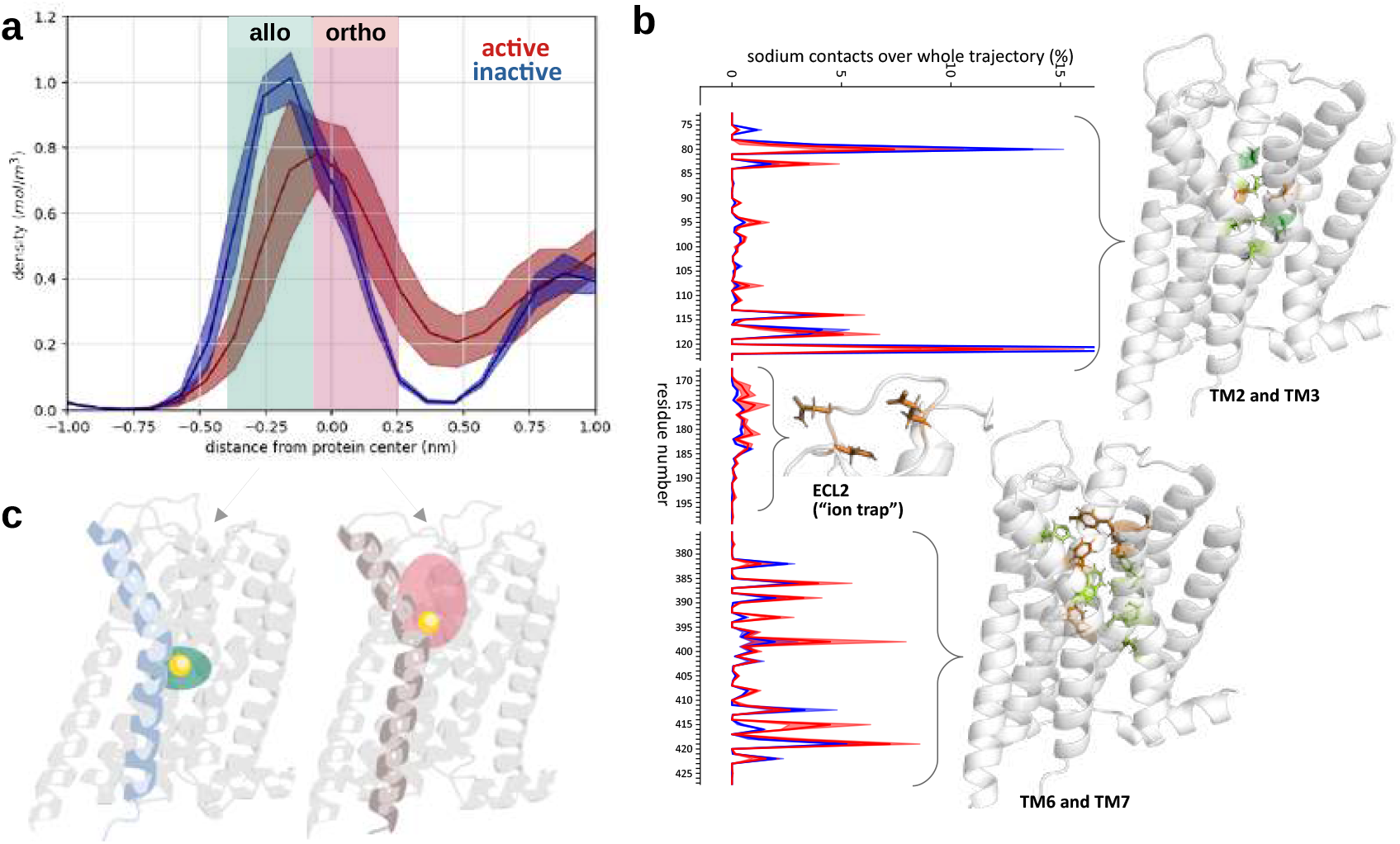
Sodium ion binding reshapes allosteric sites in DRD2. **(a)** Sodium density profiles reveal preferential localization in the allosteric pocket (-0.25 nm) in the inactive state and the orthosteric pocket in the active state. **(b)** Per-residue sodium contact frequencies identify key interaction hotspots. Known interaction sites are shown in green (dark green: D2.50 and D3.32), and novel contacts are highlighted in orange. **(c)** Schematic of the allosteric (green) and orthosteric pocket (pink).

*Na*^+^ exhibited distinct localization patterns and stably occupied the deep allosteric pocket in the inactive state, but remained closer to the orthosteric pocket and the extracellular side in the active state, suggesting different effects on the two states (Fig. 1a, c). ^5,6,23–26^

In addition to canonical binding sites, ^5–7,23–26^ we identified seven previously unreported *Na*^+^ contact residues within the transmembrane domain (V2.53_83_, C3.36_118_, F6.44_382_, F6.51_389_, Y7.35,_408_; S7.36_409_ and T7.39_412_), as well as sites in the flexible extracellular loops ^28^ acting as “ion traps”^5^ and likely guiding *Na*^+^ into the receptor (fig. 1b).

Time-resolved analysis showed rapid *Na*^+^ entry and stabilization in the allosteric pocket in the inactive state, while in the active state, *Na*^+^ entry was slower and more time was spent in the orthosteric pocket formed by D3.32_114_, C3.36_118_, W6.48_386_ and F6.52_390_ (SI section *Na*^+^ contact sites). ^5–7,23–26^ In addition, sodium ions exhibited dynamic exchange with the extracellular environment only in the active state, not in the inactive state.

## 3 *Na*^+^ triggers long-range allosteric remodeling in GPCR dynamics

It is well established that sodium ions enhance antagonist binding and stabilize the inactive state of class A GPCRs.^5–7, 23–26^ To explore sodium’s impact on receptor dynamics, we conducted simulations of four systems: active and inactive states with *Na*^+^ freely and entering the receptor, and both states with *Na*^+^ restrained outside the receptor.

Principal component analysis (PCA) of backbone dynamics revealed that sodium binding alters global conformational motions. The dominant motion (PC1) captured the opening and closing of the intracellular cleft (ICC), whereas PC2 reflected movements of the extracellular cleft and extracellular ligand-binding regions (ECC).

In *Na*^+^-bound active-state simulations, intracellular cleft closure along PC1 was observed, consistent with inactivation, while inactive-state simulations showed reduced motion and increased stability. In contrast, simulations performed in the absence of sodium revealed a stabilized active state characterized by an open intracellular cleft, supporting the notion that sodium functions as a negative allosteric modulator by favoring transitions toward the inactive state. Although increased motion along PC2 was detected in sodium-free inactive-state simulations, no transition to an active-like conformation occurred(fig. 2a).

**Figure 2:**
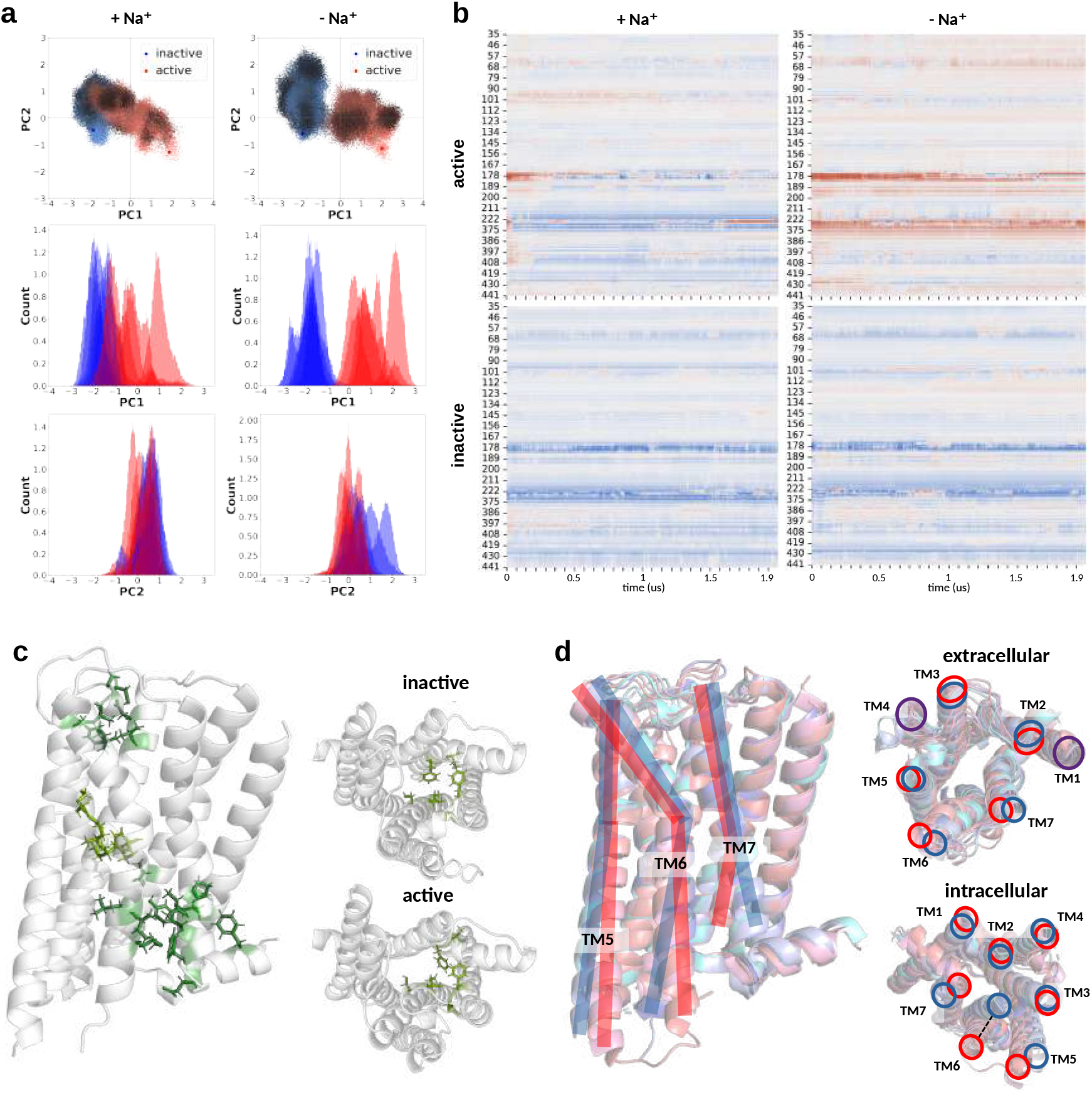
Sodium ions modulate receptor dynamics and residue-residue interactions. **(a)** Principal component analysis (PCA) reveals conformational shifts: Na^+^ promotes intracellular cleft closure in the active state (PC1), while Na^+^-free systems show enhanced extracellular flexibility (PC2). **(b)** *relDist* analysis tracks transitions toward inactive-state-like conformations, highlighting regions undergoing early conformational change. **(c)** Residue-residue contact differences across systems show reorganization of interaction networks with or without Na^+^. **(d)** Overlay of available DRD2 crystal structures illustrates key structural differences between active and inactive states.

To quantify local conformational changes, we developed a residue-level metric called relDist, which compares the position of each residue to the reference active and inactive structures (see Supplementary Information, section Analysis of conformational transitions with relDist). Positive relDist values indicate a shift toward the inactive state, whereas negative values indicate a shift toward the active state.

The relDist analysis revealed that *Na*^+^-bound active-state simulations transitioned first at the intracellular ends of TM5, TM7, and the central region of TM3, followed by TM6 and extracellular regions of TM4, TM5, TM2, TM3, and ECL1 toward an inactive-like conformation. In sodium-free active states, the receptor largely retained an active-like conformation, with shifts limited to the intracellular end of TM5 and the extracellular region of TM7, consistent with PCA results.

In the inactive state, *Na*^+^-free systems showed increased shifts toward the active state, particularly in extracellular regions of TM2 (residues 79-90), and TM6 and TM7 (residues 386-408). The central region of TM3 (residues 112-123) remained similar across states without *Na*^+^, but differed substantially in *Na*^+^-bound simulations (fig. 2b).

Comparison of *Na*^+^ position and relDist profiles revealed a correlation between the presence of *Na*^+^ in the sodium pocket and conformational changes of TM7, consistent with previous reports.^7^

These observations suggest that *Na*^+^ alters inter-residue interactions, triggering long-range allosteric changes that involve the NPxxY motif and the sodium pocket.^3,5^ Thus, *Na*^+^ binding exerts extensive allosteric effects, reshaping receptor conformation and dynamics.^5–7, 23–26^

## 4 *Na*^+^ reshapes residue interaction networks in active and inactive states

Residue-residue contact analysis revealed that *Na*^+^ stabilizes specific interhelical interactions, particularly within the extracellular and intracellular clefts. In the absence of *Na*^+^, pronounced differences between active and inactive states were observed, especially in the ICC (TM6, TM7, intracellular helix) and ECC (TM3–TM5), which are consistent with structural differences in the available experimental structures of DRD2^29–34^(fig. 2c and d, SI fig. 2).

In the *Na*^+^-bound active state, interaction patterns shifted toward those observed in the inactive state, indicating that sodium reorganizes residue interaction networks to promote inactivation. Some differences persisted at the allosteric binding site, where contacts were stronger in the inactive state, suggesting that *Na*^+^ preferentially stabilizes the ECC and central receptor regions.

Comparison with the relDist analysis showed that the central regions of TM3 retained active-like features in *Na*^+^-free systems but adopted an inactive-like conformation in *Na*^+^-bound simulations. Within this region, weaker interhelical residue-residue interactions were observed, particularly in *Na*^+^-free systems.

Notably, differences between active and inactive states were more pronounced in sodium-free systems than in sodium-bound systems, extending beyond the sodium pocket, highlighting the long-range impact of sodium on allosteric networks.^5–7, 23–26^

## 5 Free energy scan reveals residues critical for sodium ion interactions

Building on our previous work examining the effects of mutations on the active–inactive equilibrium of DRD2, ^19^ we investigated how the position of *Na*^+^ influences local state stability.

We performed an alanine scan across the receptor and calculated mutation free energy changes (ΔΔ*G*) for three sodium conditions: sodium-free, sodium in the allosteric pocket, and sodium in the orthosteric pocket (Fig.3a), using pmx.^20,21^ Positive ΔΔ*G* values indicate that the mutation stabilizes the inactive state, while negative values indicate stabilization of the active state.

**Figure 3:**
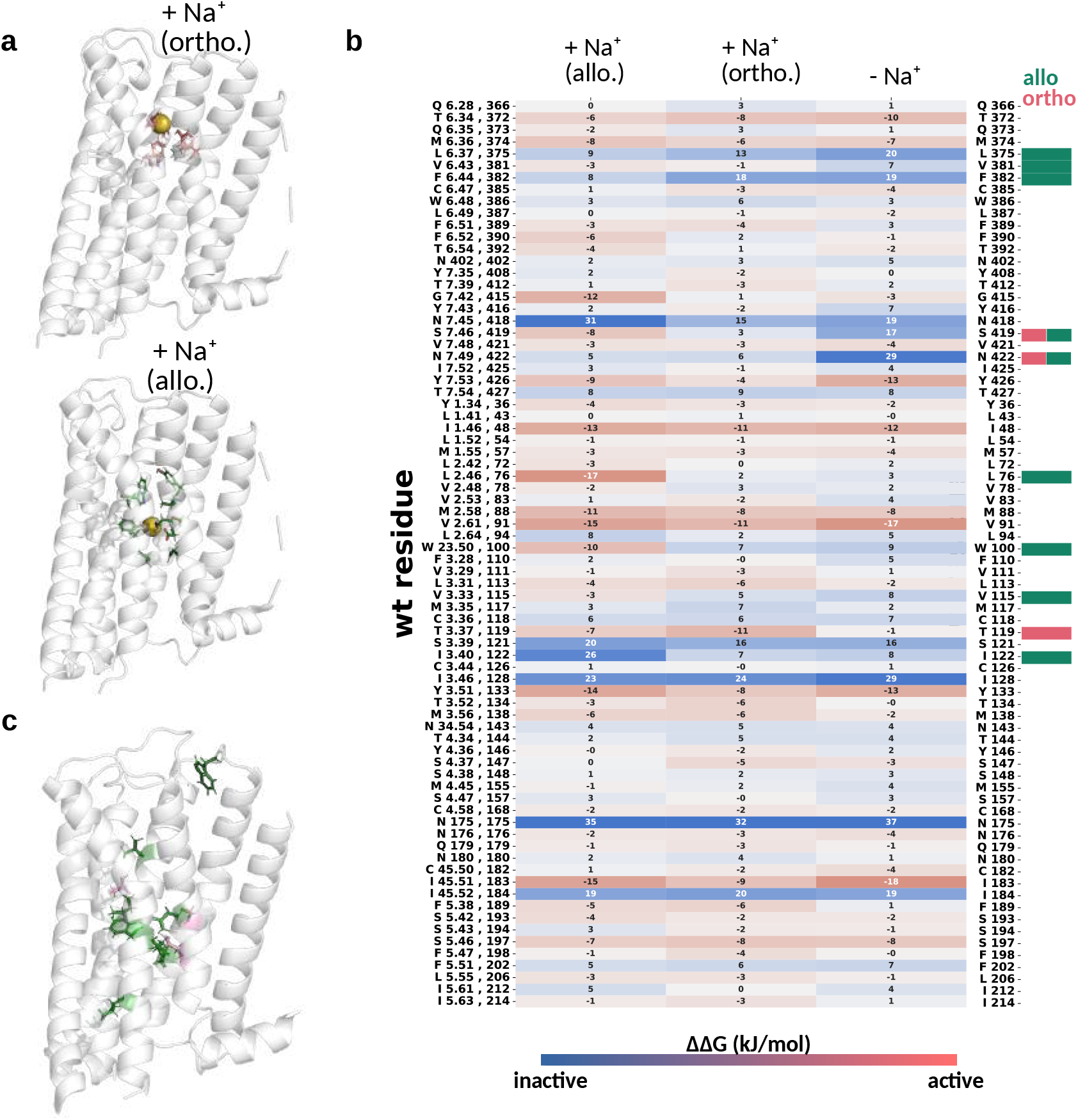
Free energy profiling identifies residues critical for Na^+^-dependent stability. **(a)** Sodium binding sites used in the free energy scans with sodium restrained at the orthosteric pocket (V2.57_87_, D3.32_114_, C3.36_118_, W6.48_386_, F6.51_389_, Y7.43_416_) and allosteric pocket (L2.46_76_, D2.50_80_, V2.53_83_, C3.36_118_, S3.39_121_, W6.48_386_, F6.44_382_, Y7.43_416_, N7.45_418_, S7.46_419_, N7.49_422_). **(b)** The ΔΔ*G* in kJ/mol of the active-inactive comparison scan. Mutants with a δ*G >* 10 for the comparison of *Na*^+^-free systems to *Na*^+^ in the orthosteric pocket are highlighted in pink and compared to *Na*^+^ in the allosteric pocket in green (second column **(b)**). **(c)** Structural mapping of Na^+^-sensitive mutations identifies key residues across active and inactive states.

To quantify sodium’s per-residue impact, we compared ΔΔ*G* values across conditions and calculated δ*G* = ΔΔΔ*G*, as the difference in ΔΔ*G* between systems with and without sodium in the allosteric pocket, which represents the non-additive free energy shift introduced by sodium.^22^ Residues with δ*G >* 10 kJ/mol were classified as sodium-coupled. As sodium-binding interactions were removed by alanine substitution, we expected a destabilization of sodium-preferred interactions.

Compared to the sodium-free system, the active state was most destabilized by mutations when *Na*^+^ was bound to the allosteric pocket, reflected by a decrease in cumulative mutation free energy ( ∑ΔΔ*G*) by −86.7 *±* 5.0 kJ/mol (orthosteric sodium) and −114.3 *±* 7.2 kJ/mol (allosteric sodium). In the inactive state, the opposite trend was observed, with increases of +22.9 *±* 5.2 kJ/mol (orthosteric sodium) and +36.5 *±* 4.9 kJ/mol (allosteric sodium) (see SI section 2).

Together, these results confirm a stronger influence of allosteric sodium on the active state, reinforcing earlier findings that highlight sodium’s state-selective allosteric role in GPCRs.^5–7, 23–26^

We identified a set of sodium-coupled residues located primarily near the allosteric (8 total: L6.37_375_A, V6.43_381_A, F6.44_382_A, N7.45_418_A, L2.46_76_A, W23.50_100_A, V3.33_115_A, I3.40_122_A) and orthosteric sodium binding pockets (T3.37_119_A). Notably, some residues, including W23.50_100_ and I3.40_122_, were distant from these sites, suggesting long-range allosteric effects (Fig. 3b, c). Particularly, N7.49_422_A and S7.46_419_A responded to sodium binding in both pockets, emphasizing their role in the conformational switch. ^7^

## 6 *Na*^+^ changes allosteric coupling networks and drives receptor inactivation

Allosteric coupling analysis revealed that *Na*^+^ alters the strength and topology of energetic coupling between key residues. The allosteric coupling between the residue pairs was quantified by analyzing the nonadditivity of the mutation free energies: (ΔΔ*G*_*A*_ (residue A) + ΔΔ*G*_*B*_ (residue B)) - ΔΔ*G*_*AB*_ (double mutation).^22^ Residue pairs with δ*G*_*AB*_ *>* 10 kJ/mol in at least two combinations were classified as part of the allosteric coupling network. We selected residues previously implicated in sodium-mediated stabilization or mutation sensitivity^19^ and performed the analysis in four systems (active/inactive ± *Na*^+^ in the allosteric site).

Seven residues formed the core coupling network across all systems, primarily involving conserved microswitches: W6.48_386_ and C6.47_385_(CWxP motif^2,23,35^), Y3.51_133_ (DRY^1,23,36,37^), I7.52_425_ (NPxxY^1–3, 23, 35–39^), F5.47_198_, M6.36_374_ (stabilizes the inactive state^19^) and M2.58_88_, reflecting their essential role in receptor state transitions.

To identify state-specific couplings, residues were classified as *inactive-specific* if they showed strong coupling (4-6 pairs) in both inactive systems but no or weak coupling (2-3 pairs) in only one active state. The inverse defined *active-specific* residues. We identified four inactive-specific residues: I45.51_183_, T6.32_374_, N7.45_418_ (sodium binding pocket^1–3, 23, 35–39^), Y7.53_426_ (NPxxY motiv), mostly involved in sodium coordination or inactive state stabilization.^19^ In addition, five active-specific residues were found: W23.50_100_, N175, F6.44_382_ (PIF motif^1, 2, 23, 35, 37^), S7.46_419_, and N7.49_422_, further supporting their role in active-state stabilization. ^19^

Active-specific residues localized mainly to the receptor core, whereas inactive-specific residues clustered near extracellular or intracellular regions.

These findings support our prior hypothesis^19^ that inactivation is associated with disrupted coupling in the receptor core, and highlight the *Na*^+^ pocket (S3.39_121_, D2.50_80_, N7.49_422_, S7.46_419_) as essential for regulating active-state stability. ^5–7, 23–26^

Sodium altered coupling networks predominantly in the active state, where we detected five residues with altered coupling patterns compared to only three in the inactive state. Notably, N7.45_418_ (inactive-specific residue) gained coupling in the sodium-bound active state, whereas S7.46_419_ (active-specific coupling) lost coupling upon sodium binding; no such transitions were observed in the inactive systems. These results suggest that *Na*^+^ shifts the active receptor toward an inactive-like coupling network, potentially initiating inactivation (Fig. 4, SI Fig. 4).

**Figure 4:**
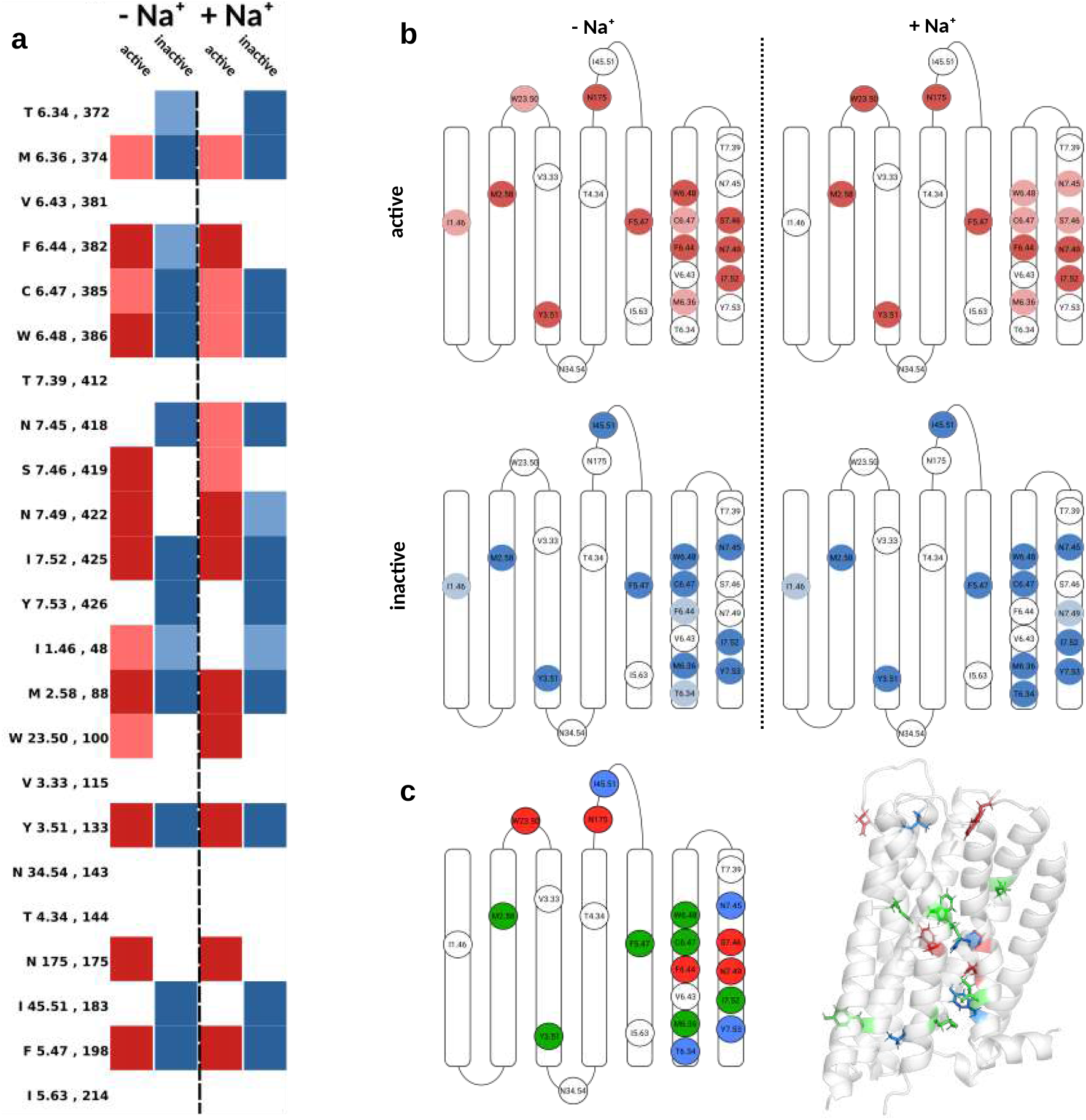
Na^+^ modulates allosteric coupling networks through inter-residue interactions. **(a)** Double mutant coupling strength matrix (*δG*) identifies residue pairs with strong non-additive allosteric coupling. Residues that are highly coupled (at least 4 coupled pairs out of 6) are shown in dark colors, those coupled to only two to three other residues shown in light colors. **(b)** Coupled residue networks mapped to a DRD2 schematic in active, inactive, Na^+^-bound, and Na^+^-free states highlight Na^+^-induced shifts in coupling topology. **(c)** Overview of state specific coupling networks. Residues that are coupled in both states are shown in green, active and inactive specific residues shown in red and blue respectively and mapped to the structure.

Furthermore, N7.45_418_, N7.49_422_, and Y7.53_426_, which form a bridge between the *Na*^+^ pocket and the NPxxY motif,^7,8^ were part of the coupling network only in sodium-bound systems (Fig.4). Because these residues are implicated in receptor-water interactions,^7,8,40^ we next examined the role of sodium in modulating hydration dynamics.

## 7 Disruption of GPCR hydration underlies sodium-mediated allosteric control

Water plays a critical role in GPCR activation.^7,8,40^ Our simulations revealed that a continuous internal water column spanning the transmembrane region formed only in the active, *Na*^+^-free state. In all other systems, a hydration discontinuity (“water gap”) appeared near N7.49_422_–Y7.53_426_.

Extracellular cavity hydration was higher in active than inactive *Na*^+^-bound systems, whereas sodium-free inactive systems showed increased hydration (Fig. 5a), consistent with enhanced flexibility observed by PCA.

**Figure 5:**
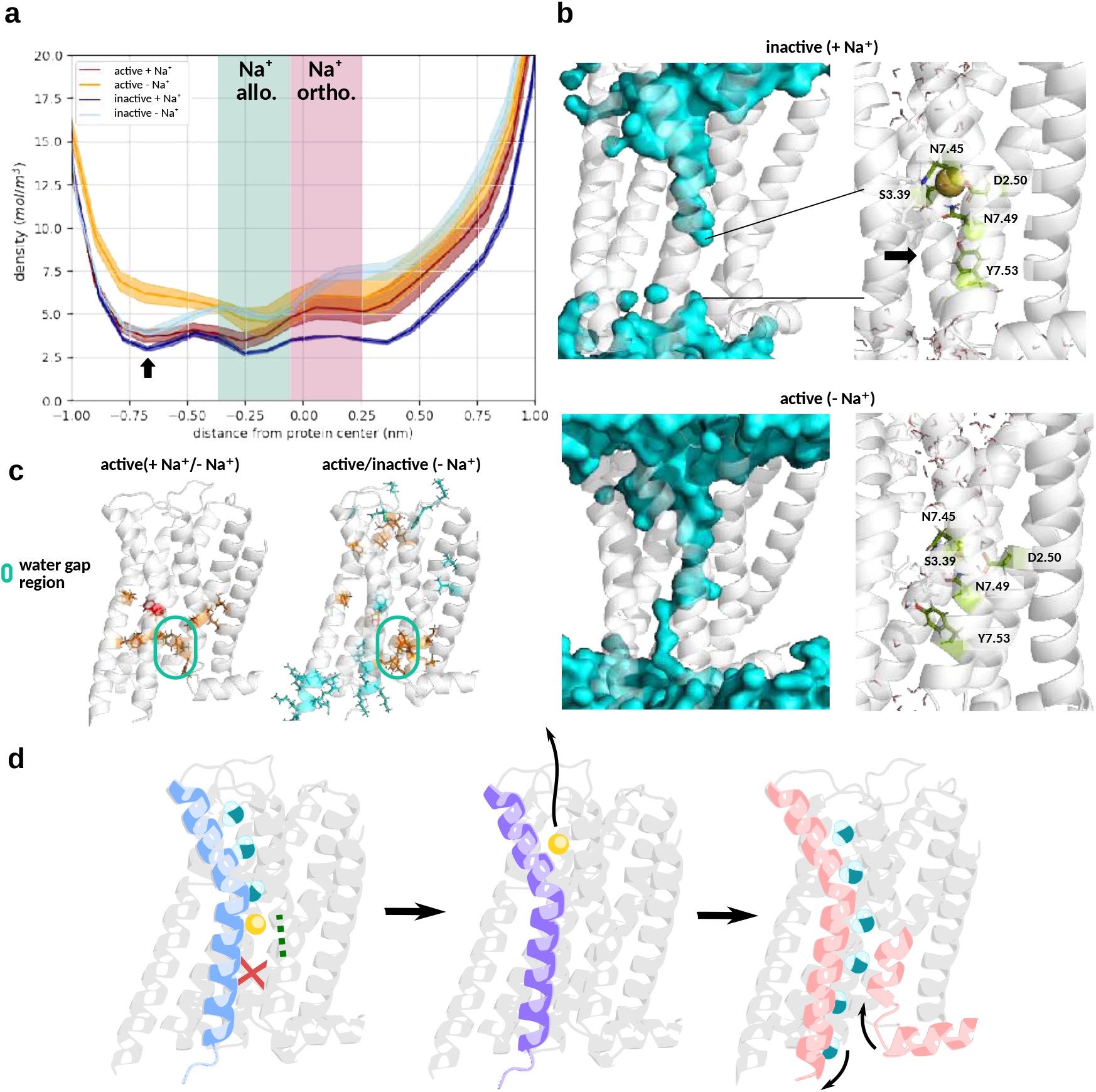
Na^+^ alters internal hydration and disrupts water channel formation. **(a)** Water density plots reveal a continuous water column only in the active Na^+^-free state, while other systems exhibit a distinct “water gap” near N7.49–Y7.53 (black arrow). **(b)** Structural representation of the water gap and hydrated channel illustrates loss of hydration continuity in Na^+^-bound systems. **(c)** Residue-wise water contact differences across systems. Water contacts enriched in active states are shown in red (Na^+^-bound) and orange (Na^+^-free); inactive-enriched contacts are shown in dark and light blue, respectively. **(d)** Schematic of the activation process. Upon Na^+^ leaving the coupling within TM7 changes (green dotted line) and finally leads to TM7 moving inward and TM6 moving outward, thereby allowing full hydration.

In the intracellular region, water density was reduced in inactive and active state sodium-bound systems, whereas a hydrated channel spanning TM2, TM3, TM6, and TM7 was preserved only in the active *Na*^+^-free systems (Fig. 5b). This matches the PCA and relDist results that identified state-dependent differences in the intracellular region. Residue-level water contact analysis showed that sodiums effects on hydration were primarily restricted to the ECC in both active and inactive systems. The water gap itself remained stable across these sodium-bound states.

Comparison of active and inactive sodium-free states revealed pronounced differences in water interactions within the gap, with increased hydration of TM3, TM5, and TM6 only in the active state. Key residues contributing to this effect included TM1: I1.46_48_, N1.50_52_; TM2: I2.43_73_; TM3: L3.43_125_, I3.46_128_, S3.47_129_; TM5: F5.51_202_, Y5.58_209_; TM6: V6.40_378_; and TM7: A7.47_420_, V7.48_421_, Y7.53_426_, F7.56_429_. In active-state comparisons, significant sodium-induced hydration changes were confined to the water gap region (Fig. 5 c, SI Fig. 5).

This water gap coincides with the segment of TM7 that adopts an inactive-like conformation in sodium-bound active states. The region hosts a cluster of water molecules adjacent to the sodium-binding pocket and the NPxxY motif (N7.45_418_, N7.49_422_, S3.39_121_, D2.50_80_).^7,8^ Upon sodium binding to the deep allosteric pocket, local hydrogen bonding is perturbed: N7.49_422_ is pushed outward, and Y7.53_426_ rotates upward, breaking the water chain. In the absence of *Na*^+^, N7.45_418_ and N7.49_422_ participate in water-mediated bonding that draws TM7 inward, allowing Y7.53_426_ to rotate into the receptor core. This rearrangement disrupts a hydrophobic triad formed by TM2, TM3, and TM6, thereby enabling full channel hydration.

Consistent with these observations, the allosteric coupling network analysis showed that N7.45_418_ and N7.49_422_ are only part of the allosteric coupling network in both states in presence of sodium. This reinforces their role in linking the NPxxY motif to the *Na*^+^ pocket and mediating intracellular cleft closure during sodium-induced inactivation.

## 8 Conclusion

Our findings reveal a multifaceted role of sodium ions in regulating GPCR activation. Beyond their known role in stabilizing inactive states, we show how *Na*^+^ acts as an allosteric modulator, reshaping both structural dynamics and inter-residue coupling networks in the dopamine D2 receptor.

In the active state, sodium binding initiates a cascade of conformational and interaction changes: coupling patterns shift toward inactive-like configurations, the water pathway is disrupted, and key helices undergo structural rearrangement. These shifts were quantified using PCA, relDist metrics, and mutation-induced free energy changes, all pointing to sodium-driven inactivation.

Importantly, we demonstrate that *Na*^+^ binding alters long-range coupling beyond its immediate pocket. Residues several angstroms away, including those in the receptor core and intracellular domains, were functionally affected, revealing a mechanistic bridge between local binding and global conformational control.

The hydration gap observed only in *Na*^+^-bound systems introduces a new layer of regulation, since water exclusion from critical microswitch regions such as *Na*^+^ pocket and the NPxxY motif appears to prevent activation. ^7^ This supports recent theories proposing water as an integral player in GPCR signal transduction, alongside ions and ligands.^41,42^

Taken together, our results suggest that sodium regulates GPCR activity via a triad of mechanisms: stabilizing inactive-specific contact networks, modulating long-range allosteric coupling, and disrupting the water-mediated activation channel. Given the high sequence and structural conservation of the *Na*^+^-binding site and associated microswitches across class A GPCRs,^1–3, 19, 23, 35–39^ we propose that the sodium-induced coupling network remodeling and hydration gap disruption observed in DRD2 likely represent a conserved regulatory mechanism among class A GPCRs.

These insights offer new directions for drug design. Antagonists may benefit from mimicking sodium’s effects by promoting binding of *Na*^+^ or allosterically stabilizing inactive-like networks.^43^ In contrast, agonists could be optimized to replace *Na*^+^ in the allosteric pocket or restore hydration continuity, thereby enhancing receptor activation.

## 9 Methods

### 9.1 Structure selection and system setup

Structures were chosen and prepaired as described by Schmidt and de Groot.^19^ The highest-resolution structures representing the inactive (6cm4, 2.9 Å)^31^ and active (7jvr, 2.8 Å)^29^ conformational states were chosen and oriented in a lipid bilayer using the OPM server. ^44^

Non-native mutations present in 6cm4 (A122I, A375L, A379L) were reverted to wild-type residues in PyMOL. The unresolved loop (residues 139–144) was modeled based on the corresponding region in 7jvr, while unresolved termini were excluded. The intracellular loop 3 (ICL3; V222–Q364), which is unresolved in both structures, was omitted, effectively dividing the receptor into two subunits: S1 (TM1–5, N35–R221) and S2 (TM6–7, Q354–L441). Both termini of each subunit were capped with ACE and CT3 groups using CHARMM-GUI.^45^ System setup in CHARMM-GUI included embedding the receptor in a POPC lipid bilayer, solvating with TIP3P water, and adding 150 mM NaCl. ^46,47^ All simulations used the CHARMM36m force field.^48^

### 9.2 System Equilibration and Simulation

Simulations were conducted with GROMACS 2022.3^49–51^ using the CHARMM36m force field^48^ and TIP3P water model^52^ at 310.5K. Energy minimization was carried out using the steepest descent algorithm until convergence, followed by a six step equilibration with gradually decreasing restraints on atom positions and dihedrals. Short-range electrostatics and van der Waals interactions used a 1.2nm cutoff; long-range electrostatics were treated with PME.^53^ Force-switch (vdW) and potential-shift (Coulomb) modifiers ensured smooth transitions. Covalent bonds involving hydrogen were constrained with LINCS, ^54^ using a 2 fs timestep.

Initial NVT equilibration used the Berendsen thermostat (*τ* = 1 ps), followed by four NPT steps with Berendsen thermostat/barostat (*τ* = 5 ps, semi-isotropic). Production runs (600ns, NPT) used a Nose–Hoover thermostat (*τ* = 1 ps) and semi-isotropic Parrinello–Rahman barostat (*τ* = 5 ps). Where indicated, position restraints were applied to all protein backbone *C*_*α*_ atoms and *Na*^+^ ions with a force constant of 1000 kJ mol^−1^ nm^−2^.

Each system was run for 2 *µ*s and 5 replicas.

### 9.3 Setup for Free Energy Alchemical Transitions with PMX

Alchemical transitions were prepared with PMX,^20,21^ following the protocol of Aldeghi et al.^55–57^ Mutations, hybrid structures, and topologies were generated accordingly. Mutant systems were energy-minimized and equilibrated for 500 ps at 310.5 K and 1 bar using a Langevin thermostat^58^ and Berendsen barostat, followed by a 50 ns equilibrium simulation with a Langevin thermostat and Parrinello–Rahman barostat.

From the final 20 ns of equilibration, 200 configurations were extracted at 100 ps intervals as starting points for 100 ps nonequilibrium transitions. Soft-core potentials^59^ were applied to non-bonded interactions. Long-range electrostatics were treated with PME, and short-range interactions used a 1.0 nm cutoff. All covalent bonds involving hydrogens were constrained with LINCS (time step = 2 fs).

To prevent transitions between conformational states, position restraints (1000 kJ mol^−1^ nm^−2^) were applied to all *C*_*α*_ atoms and *Na*^+^ ions throughout the simulation. *Na*^+^ ions were restrained outside the receptor to prevent interaction.

### 9.4 Calculation and Analysis of Relative Free Energy Changes and Uncertainities

Work values from nonequilibrium transitions were extracted using PMX,^21,56,57^ and free energy differences for active and inactive states were computed via the Bennett Acceptance Ratio (BAR) method. Uncertainties of the Δ*G* were obtained via bootstrapping over the combined work value distribution of triplicates. The resulting values, 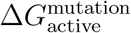 and 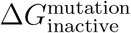, were used to compute the relative free energy change between states. The shift in conformational equilibrium upon mutation was quantified as:

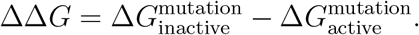

The final uncertainties of the relative free energy change ΔΔ*G* were calculated via error propagation from the Δ*G* uncertainties.

## 10 Acknowledgement

Funding from the Max Planck Society is gratefully acknowledged. LS is funded by the IMPRS for Physics of Biological and Complex Systems Excellence Fellowship.

## 11 Author Contribution

Conceptualization: LS, BLG Methodology: LS, BLG Investigation: LS Visualization: LS Supervision: BLG Writing—original draft: LS Writing—review and editing: BLG, LS

## 13 Data and materials availability

Additional information can be found in the Supplementary Information.

## References

(1) Latorraca, N. R.; Venkatakrishnan, A. J.; Dror, R. O. GPCR Dynamics: Structures in Motion. Chemical Reviews 2017, 117, 139–155.

(2) Zhou, Q. et al. Common activation mechanism of class A GPCRs. eLife 2019, 8, e50279.

(3) Katritch, V.; Cherezov, V.; Stevens, R. C. Structure-Function of the G Protein–Coupled Receptor Superfamily. Annual Review of Pharmacology and Toxicology 2013, 53, 531– 556.

(4) Agasid, M. T.; Sørensen, L.; Urner, L. H.; Yan, J.; Robinson, C. V. The Effects of Sodium Ions on Ligand Binding and Conformational States of G Protein-Coupled Re-ceptors—Insights from Mass Spectrometry. Journal of the American Chemical Society 2021, 143, 4085–4089.

(5) Selent, J.; Sanz, F.; Pastor, M.; De Fabritiis, G. Induced Effects of Sodium Ions on Dopaminergic G-Protein Coupled Receptors. PLOS Computational Biology 2010, 6, 1–6.

(6) Michino, M.; Free, R. B.; Doyle, T. B.; Sibley, D. R.; Shi, L. Structural basis for Na+sensitivity in dopamine D2 and D3 receptors. Chemical Communications 2015, 51, 8618–8621.

(7) Katritch, V.; Fenalti, G.; Abola, E. E.; Roth, B. L.; Cherezov, V.; Stevens, R. C. Allosteric sodium in class A GPCR signaling. Trends in Biochemical Sciences 2014, 39, 233–244.

(8) White, K. L.; Eddy, M. T.; Gao, Z.-G.; Han, G. W.; Lian, T.; Deary, A.; Patel, N.; Jacobson, K. A.; Katritch, V.; Stevens, R. C. Structural Connection between Activation Microswitch and Allosteric Sodium Site in GPCR Signaling. Structure 2018, 26, 259– 269.e5.

(9) Venkatakrishnan, A. J.; Ma, A. K.; Fonseca, R.; Latorraca, N. R.; Kelly, B.; Betz, R. M.; Asawa, C.; Kobilka, B. K.; Dror, R. O. Diverse GPCRs exhibit conserved water networks for stabilization and activation. Proceedings of the National Academy of Sciences 2019, 116, 3288–3293.

(10) Radoux-Mergault, A.; Oberhauser, L.; Aureli, S.; Gervasio, F. L.; Stoeber, M. Subcellular location defines GPCR signal transduction. Science Advances 2023, 9, eadf6059.

(11) de Felice, A.; Aureli, S.; Limongelli, V. Drug Repurposing on G Protein-Coupled Receptors Using a Computational Profiling Approach. Frontiers in Molecular Biosciences 2021, 8 .

(12) Hedderich, J. B.; Persechino, M.; Becker, K.; Heydenreich, F. M.; Gutermuth, T.; Bouvier, M.; Bünemann, M.; Kolb, P. The pocketome of G-protein-coupled receptors reveals previously untargeted allosteric sites. Nature Communications 2022, 13, 2567.

(13) Limongelli, V. Ligand binding free energy and kinetics calculation in 2020. WIREs Computational Molecular Science 2020, 10, e1455.

(14) Yuan, X.; Raniolo, S.; Limongelli, V.; Xu, Y. The Molecular Mechanism Underlying Ligand Binding to the Membrane-Embedded Site of a G-Protein-Coupled Receptor. Journal of Chemical Theory and Computation 2018, 14, 2761–2770.

(15) Dror, R. O.; Arlow, D. H.; Maragakis, P.; Mildorf, T. J.; Pan, A. C.; Xu, H.; Borhani, D. W.; Shaw, D. E. Activation mechanism of the β2-adrenergic receptor. Proceedings of the National Academy of Sciences 2011, 108, 18684–18689.

(16) Xu, P. et al. Structural insights into the lipid and ligand regulation of serotonin receptors. Nature 2021, 592, 469–473.

(17) Jefferson, R. E.; Oggier, A.; Füglistaler, A.; Camviel, N.; Hijazi, M.; Villarreal, A. R.; Arber, C.; Barth, P. Computational design of dynamic receptor—peptide signaling complexes applied to chemotaxis. Nature Communications 2023, 14, 2875.

(18) Aranda-García, D. et al. Large scale investigation of GPCR molecular dynamics data uncovers allosteric sites and lateral gateways. Nature Communications 2025, 16, 2020.

(19) Schmidt, L.; de Groot, B. Identification of allosteric sites and ligand-induced modulation in the dopamine receptor through large-scale alchemical mutation scan. Chemical Science 2025, –.

(20) Seeliger, D.; de Groot, B. L. Protein Thermostability Calculations Using Alchemical Free Energy Simulations. Biophysical Journal 2010, 98, 2309–2316.

(21) Gapsys, V.; Michielssens, S.; Seeliger, D.; de Groot, B. L. pmx: Automated protein structure and topology generation for alchemical perturbations. Journal of Computational Chemistry 2015, 36, 348–354.

(22) Werner, M.; Gapsys, V.; de Groot, B. L. One Plus One Makes Three: Triangular Coupling of Correlated Amino Acid Mutations. The Journal of Physical Chemistry Letters 2021, 12, 3195–3201, PMID: 33760609.

(23) Yang, D.; Zhou, Q.; Labroska, V.; Qin, S.; Darbalaei, S.; Wu, Y.; Yuliantie, E.; Xie, L.; Tao, H.; Cheng, J.; Liu, Q.; Zhao, S.; Shui, W.; Jiang, Y.; Wang, M.-W. G proteincoupled receptors: structureand function-based drug discovery. Signal Transduction and Targeted Therapy 2021, 2059–3635.

(24) Chan, H. C. S.; Xu, Y.; Tan, L.; Vogel, H.; Cheng, J.; Wu, D.; Yuan, S. Enhancing the Signaling of GPCRs via Orthosteric Ions. ACS Central Science 2020, 6, 274–282.

(25) Draper-Joyce, C. J.; Verma, R. K.; Michino, M.; Shonberg, J.; Kopinathan, A.; Klein Herenbrink, C.; Scammells, P. J.; Capuano, B.; Abramyan, A. M.; Thal, D. M.; Javitch, J. A.; Christopoulos, A.; Shi, L.; Lane, J. R. The action of a negative allosteric modulator at the dopamine D2 receptor is dependent upon sodium ions. Scientific Reports 2018, 1208 .

(26) Zou, R.; Wang, X.; Li, S.; Chan, H. C. S.; Vogel, H.; Yuan, S. The role of metal ions in G protein-coupled receptor signalling and drug discovery. WIREs Computational Molecular Science 2022, 12, e1565.

(27) Neve, K. A.; Cumbay, M. G.; Thompson, K. R.; Yang, R.; Buck, D. C.; Watts, V. J.; DuRand, C. J.; Teeter, M. M. Modeling and Mutational Analysis of a Putative Sodium-Binding Pocket on the Dopamine D2 Receptor. Molecular Pharmacology 2001, 60, 373–381.

(28) Nicoli, A.; Dunkel, A.; Giorgino, T.; de Graaf, C.; Di Pizio, A. Classification Model for the Second Extracellular Loop of Class A GPCRs. Journal of Chemical Information and Modeling 2022, 62, 511–522, PMID: 35113559.

(29) Zhuang, Y. et al. Structural insights into the human D1 and D2 dopamine receptor signaling complexes. Cell 2021, 184, 931–942.e18.

(30) Im, D. et al. Structure of the dopamine D2 receptor in complex with the antipsychotic drug spiperone. Nature Communications 2020, 11, 6442.

(31) Wang, S.; Che, T.; Levit, A.; Shoichet, B. K.; Wacker, D.; Roth, B. L. Structure of the D2 dopamine receptor bound to the atypical antipsychotic drug risperidone. Nature 2018, 555 .

(32) Fan, L.; Tan, L.; Chen, Z.; Qi, J.; Nie, F.; Luo, Z.; Cheng, J.; Wang, S. Haloperidol bound D2 dopamine receptor structure inspired the discovery of subtype selective ligands. Nature Communications 2020, 11, 1074.

(33) Yin, J.; Chen, K.-Y. M.; Clark, M. J.; Hijazi, M.; Kumari, P.; Bai, X.-c.; Sunahara, R. K.; Barth, P.; Rosenbaum, D. M. Structure of a D2 dopamine receptor–Gprotein complex in a lipid membrane. Nature 2020, 584, 125–129.

(34) Xu, P. et al. Structural genomics of the human dopamine receptor system. Cell Research 2023, 33, 604–616.

(35) Hauser, A. S.; Kooistra, A. J.; Munk, C.; Heydenreich, F. M.; Veprintsev, D. B.; Bouvier, M.; Babu, M. M.; Gloriam, D. E. GPCR activation mechanisms across classes and macro/microscales. Nature Structural Molecular Biology 2021, 28, 879–888.

(36) Nygaard, R. et al. The Dynamic Process of 2-Adrenergic Receptor Activation. Cell 2013, 152, 532–542.

(37) Chen, K.-Y. M.; Keri, D.; Barth, P. Computational design of G Protein-Coupled Receptor allosteric signal transductions. Nature Chemical Biology 2020, 16, 77–86.

(38) Kling, R. C.; Tschammer, N.; Lanig, H.; Clark, T.; Gmeiner, P. Active-State Model of a Dopamine D2 Receptor Gi Complex Stabilized by Aripiprazole-Type Partial Agonists. PLoS ONE 2014, 9, e100069.

(39) Mafi, A.; Kim, S.-K.; Goddard, W. A. The mechanism for ligand activation of the GPCR–G protein complex. Proceedings of the National Academy of Sciences 2022, 119, e2110085119.

(40) Thomson, N. J.; Vickery, O. N.; Ives, C. M.; Zachariae, U. Ion-water coupling controls class A GPCR signal transduction pathways. bioRxiv 2021,

(41) Fried, S. D. E.; Hewage, K. S. K.; Eitel, A. R.; Struts, A. V.; Weerasinghe, N.; Perera, S. M. D. C.; Brown, M. F. Hydration-mediated G-protein-coupled receptor activation. Proceedings of the National Academy of Sciences of the United States of America 2022, 21 .

(42) Struts, A. V.; Barmasov, A. V.; Fried, S. D.; Hewage, K. S.; Perera, S. M.; Brown, M. F. Osmotic stress studies of G-protein-coupled receptor rhodopsin activation. Biophysical Chemistry 2024, 304, 107112.

(43) Zhang, M.; Chen, T.; Lu, X.; Lan, X.; Chen, Z.; Lu, S. G protein-coupled receptors (GPCRs): advances in structures, mechanisms and drug discovery. Signal Transduction and Targeted Therapy 2024, 9, 88.

(44) Pogozheva, I. D.; Joo, H.; Mosberg, H. I.; Lomize, A. L. OPM database and PPM web server: resources for positioning of proteins in membranes. Nucleic acids research 2011,

(45) Jo, S.; Kim, T.; Iyer, V. G.; Im, W. CHARMM-GUI: A web-based graphical user interface for CHARMM. Journal of Computational Chemistry 2008, 29, 1859–1865.

(46) Wu, E. L.; Cheng, X.; Jo, S.; Rui, H.; Song, K. C.; Dávila-Contreras, E. M.; Qi, Y.; Lee, J.; Monje-Galvan, V.; Venable, R. M.; Klauda, J. B.; Im, W. CHARMM-GUI Membrane Builder toward realistic biological membrane simulations. Journal of Computational Chemistry 2014, 35, 1997–2004.

(47) Lee, J. et al. CHARMM-GUI Input Generator for NAMD, GROMACS, AMBER, OpenMM, and CHARMM/OpenMM Simulations Using the CHARMM36 Additive Force Field. Journal of Chemical Theory and Computation 2016, 12, 405–413.

(48) Huang, J.; Rauscher, S.; Nawrocki, G.; Ran, T.; Feig, M.; de Groot, B. L.; Grubmüller, H.; MacKerell, A. D. CHARMM36m: an improved force field for folded and intrinsically disordered proteins. Nature Methods 2017, 14 .

(49) Bauer, P.; Hess, B.; Lindahl, E. GROMACS 2022.3 Manual. 2022; 10.5281/zenodo.7037337.

(50) Lindahl, E.; Hess, B.; van der Spoel, D. GROMACS 3.0: a package for molecular simulation and trajectory analysis. 2001, 7 .

(51) Van Der Spoel, D.; Lindahl, E.; Hess, B.; Groenhof, G.; Mark, A. E.; Berendsen, H. J. C. GROMACS: Fast, flexible, and free. Journal of Computational Chemistry 2005, 26, 1701–1718.

(52) Jorgensen, W. L.; Chandrasekhar, J.; Madura, J. D.; Impey, R. W.; Klein, M. L. Comparison of simple potential functions for simulating liquid water. The Journal of Chemical Physics 1983, 79, 926–935.

(53) Darden, T. A.; York, D. M.; Pedersen, L. G. Particle mesh Ewald: An Nlog(N) method for Ewald sums in large systems. Journal of Chemical Physics 1993, 98, 10089–10092.

(54) Hess, B.; Bekker, H.; Berendsen, H. J. C.; Fraaije, J. G. E. M. LINCS: A linear constraint solver for molecular simulations. Journal of Computational Chemistry 1997, 18, 1463–1472.

(55) Aldeghi, M.; de Groot, B. L.; Gapsys, V. In Computational Methods in Protein Evolution; Sikosek, T., Ed.; Springer New York: New York, NY, 2019; pp 19–47.

(56) Schmidt, L.; Wilson, C. J.; Behera, S.; de Groot, B. Free energy calculations for protein design. ChemRxiv 2025, –.

(57) Behera, S.; Wilson, C. J.; Schmidt, L.; de Groot, B. Free Energy Simulations to Quantitatively Study Biomolecule Stability and Binding. ChemRxiv 2025, –.

(58) Goga, N.; Rzepiela, A. J.; de Vries, A. H.; Marrink, S. J.; Berendsen, H. J. C. Efficient Algorithms for Langevin and DPD Dynamics. Journal of Chemical Theory and Computation 2012, 8, 3637–3649.

(59) Beutler, T. C.; Mark, A. E.; van Schaik, R. C.; Gerber, P. R.; van Gunsteren, W. F. Avoiding singularities and numerical instabilities in free energy calculations based on molecular simulations. Chemical Physics Letters 1994, 222, 529–539.

