## Supplementary Information for "Sodium Ions Regulate GPCR Activation by Remodeling Allosteric Coupling Networks and Hydration Patterns"

#### 1 $Na^+$ contact sites

In both states, we observed strong $Na^+$ interactions in TM2 at D2.50$_{80}$[1] and V2.53$_{83}$ for both states. In the inactive state, additional contacts were observed at A2.49$_{79}$[2] and L2.46$_{76}$[2] deeper in the cavity. While V2.53$_{83}$ showed similar interactions in both states, all other contacts in TM2, especially D2.50$_{80}$ were stronger in the inactive state.

In TM3 D3.32$_{114}$;[1,3] C3.36$_{118}$ and S3.39$_{121}$[1,3,4] were the main residues that formed $Na^+$ contacts in both active and inactive states.

Additional contacts at M3.35$_{117}$ were primarily observed in the inactive state as reported by Michino and coworkers.[2] Comparing both states, C3.38$_{118}$ shows slightly more contacts in the active state simulations, whereas contacts to M3.35$_{117}$ were more prevalent in the inactive state. Notably, S3.39$_{121}$ exhibited stronger $Na^+$ interactions in the inactive state simulations.

In TM6, W6.48$_{386}$[1] and F6.51$_{389}$ formed similarly strong $Na^+$ contacts in bth states. An additional contact at F6.44$_{382}$ was stronger in the inactive state. This residue is located deeper in the cavity, below the W6.48$_{386}$ toggle switch at the allosteric Na binding site.[1] Additional short-lived contacts, primarily observed in the active state above the W-toggle switch at $H6.55_{393}$[3] further indicate, that $Na^+$ spends more time in the orthosteric binding site in the active state receptor.

In TM7, $Na^+$ contact sites common to both states included $Y7.35_{408}$, $S7.36_{409}$ at the extracellular top of TM7, $T7.39_{412}$ and $G7.42_{415}$;[1] $Y7.43_{416}$;[1] $N7.45_{418}$;[1] $S7.46_{419}$[1] near the orthosteric binding site. An additional contact at $N7.49_{422}$,[5] was detected deeper in the cavity at the allosteric binding site. While $G7.42_{415}$ and $Y7.43_{416}$ $Na^+$ contacts were predominantly observed in the active state and $T7.39_{412}$ showed stronger interactions in the inactive state, most TM7 interaction sides were present in both states.

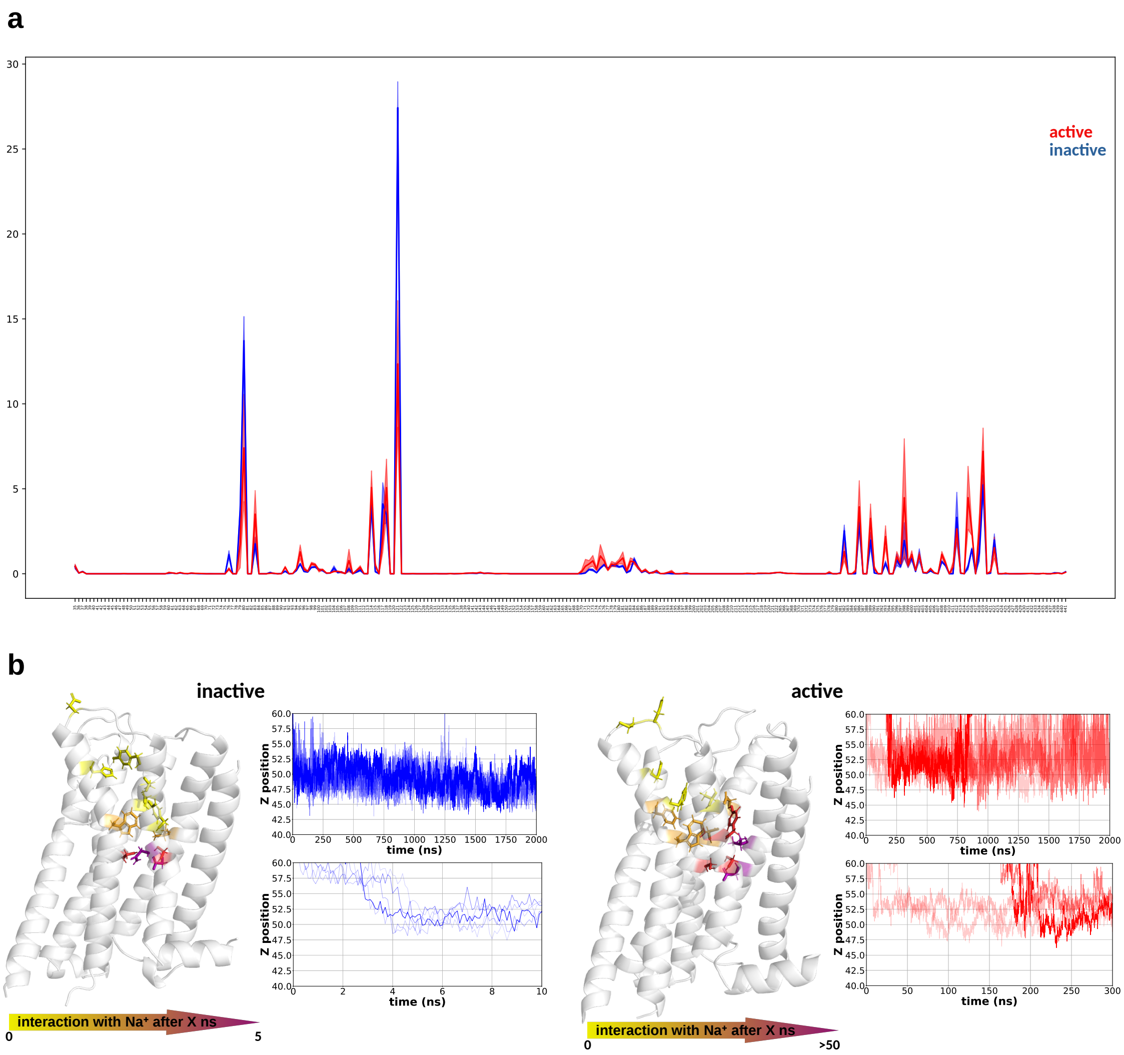


Figure 1: **Sodium contacts and entry pathway.** **(a)** Per-residue sodium contact frequencies. **(b)** Time-resolved $Na^+$ entry pathways show distinct binding dynamics. In the inactive state, $Na^+$ rapidly occupies the allosteric pocket, while in the active state it first localizes to the orthosteric region before entering deeper pockets.

#### 2 Analysis of conformational transitions with relDist

For each simulation frame, we computed the similarity of the receptor's backbone (C$\alpha$ atoms) to both reference states (active and inactive). Specifically:

1. Global Alignment: Each simulation frame was structurally aligned to the active and

inactive crystal structures to remove overall rotation/translation effects.

2. C$\alpha$ Distance Calculation: For every residue $i$, we measured the Euclidean distance between its C$\alpha$ position in the simulation frame to both the active and inactive reference structures.

$$d_{i,\text{sim}-\text{active}}, \quad d_{i,\text{sim}-\text{inactive}}$$

3. Relative Distance (relDist): We subtracted the distances from step 2 to the active state from the distance to the inactive state for each residue:

$$\text{relDist}_i = d_{i,\text{sim}-\text{inactive}} - d_{i,\text{sim}-\text{active}}$$

A positive $\text{relDist}_i$ value indicates that residue $i$ in the simulation frame is closer to the inactive conformation, while a negative value indicates a shift toward the active state.

#### 3 Inter-residue contacts

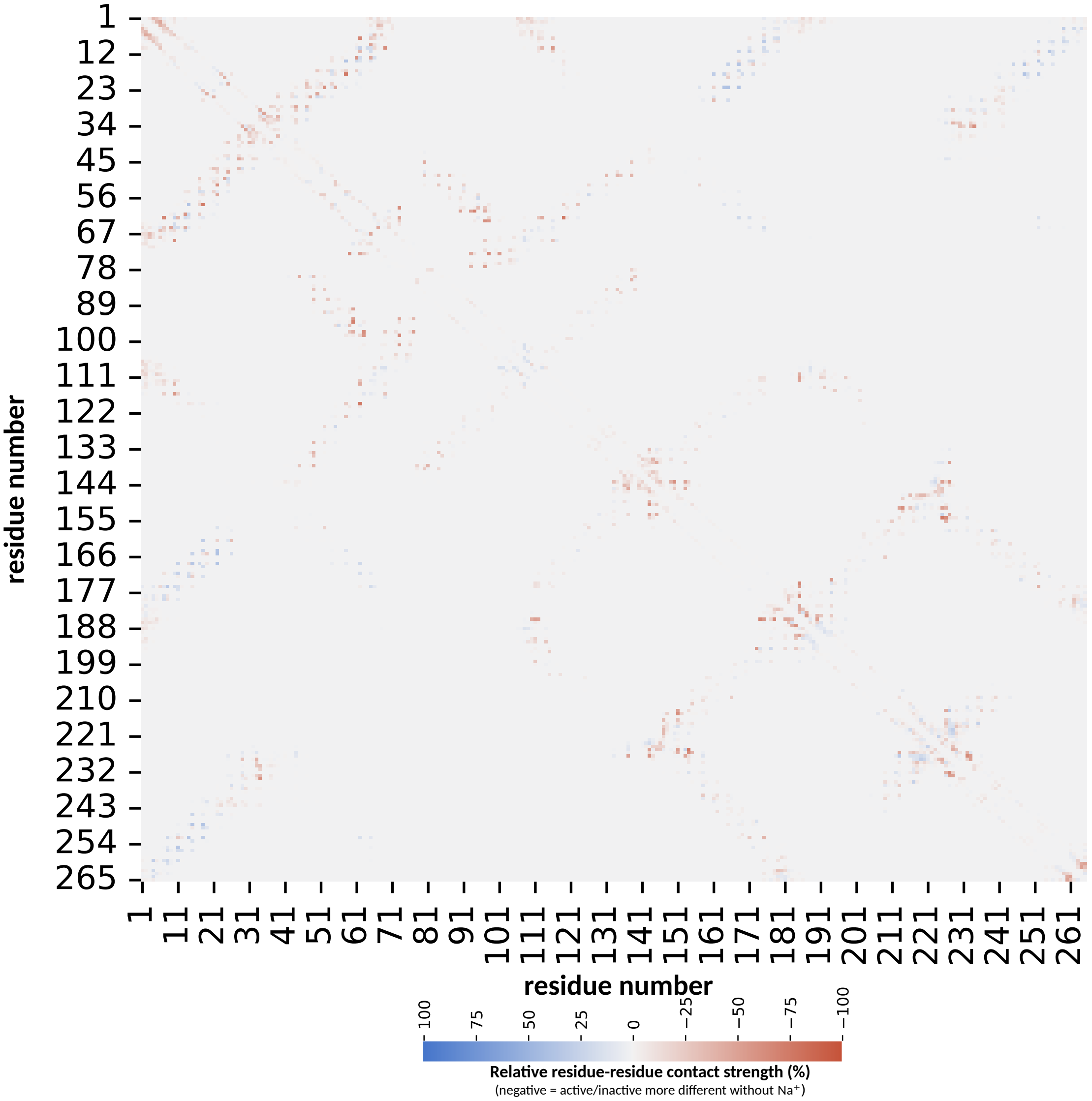


Figure 2: **Relative residue–residue interaction differences.** The contact plot compares interactions that differ most strongly between active and inactive states for sodium-free and sodium-interacting systems. More negative values indicate greater differences between active and inactive states in sodium-free simulations.

### 4 Single mutation free energy scan with sodium at different positions

This scan quantifies the effect of mutation on the active and inactive systems separately, using the thermodynamic cycle in figure 3c. Positive values indicate a destabilizing effects, whereas negative values indicate stabilizing effects of the mutation.

Sodium-coupled residues differed between receptor states. In the inactive state, only five residues (I3.40$_{122}$A, L6.41$_{379}$A, F6.44$_{382}$A, W6.48$_{386}$A, N7.49$_{422}$A) were significantly affected. In contrast, the active state showed stronger sodium effects, with seven residues exhibiting substantial free energy shifts (L2.46$_{76}$A, W23.50$_{100}$A, S3.39$_{121}$A, W6.48$_{386}$A, N7.45$_{418}$A, S7.46$_{419}$A, N7.49$_{422}$A), particularly when sodium was bound to the allosteric pocket (SI Fig. 2). This further highlights the stronger effect of $Na^+$ on the active state.

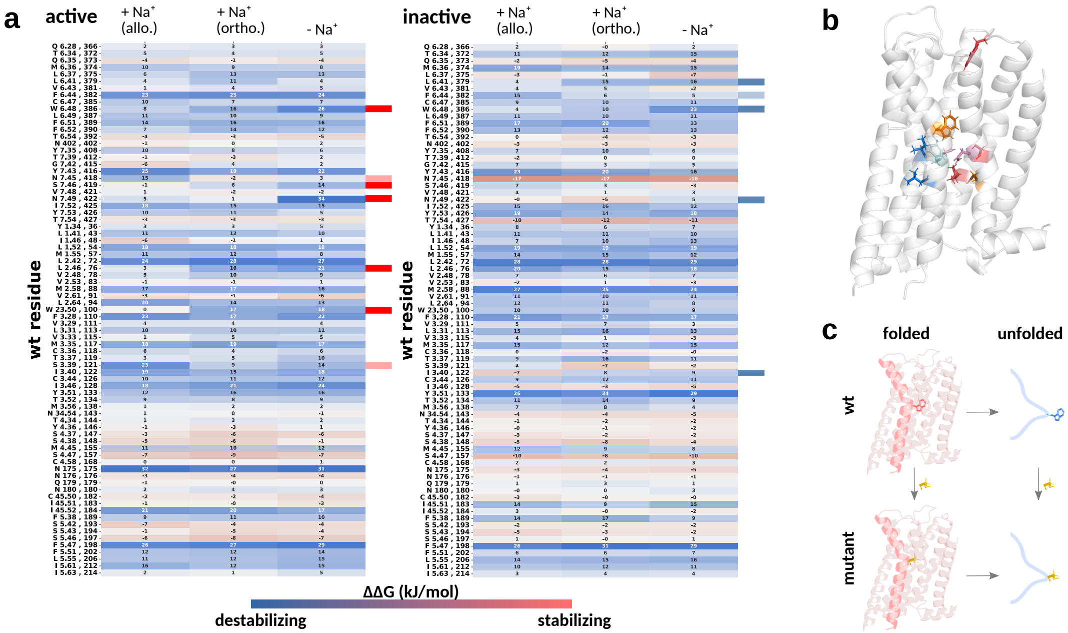


Figure 3: **Free energy single mutation scan for active and inactive state separately.** Mutants that are destabilized by $Na^+$ are marked with light red/blue color, mutants stabilized by $Na^+$ marked in dark red/blue (, 4th column a inactive/active). B shows the marked residues on the structure, the colors correspond to the marker colors, except for residues that behave the same in active and inactive are colored orange. C shows an example thermodynamic cycle used for the calculation.

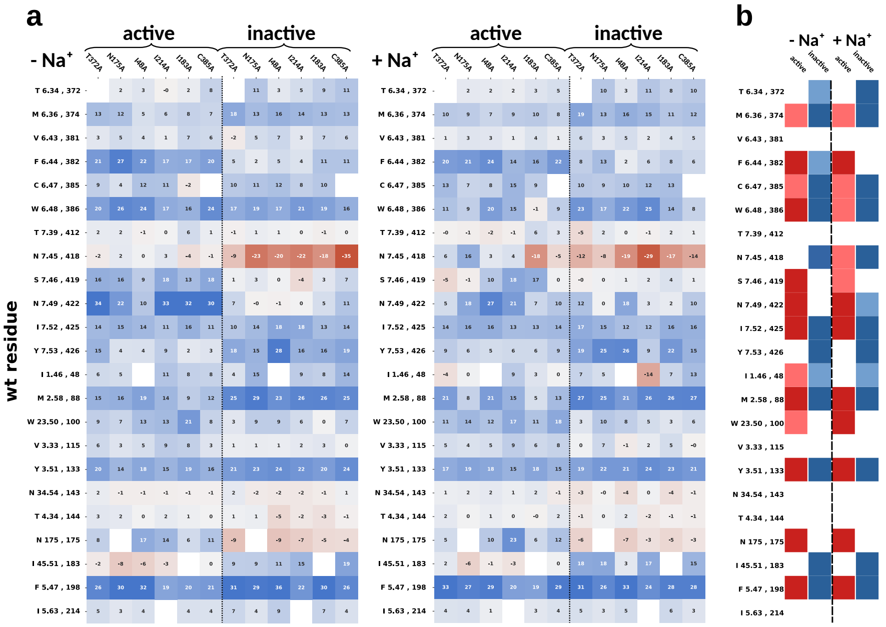

Figure 4: **Free energy double mutation scan for $Na^{+}$-bound and $Na^{+}$-free systems.**

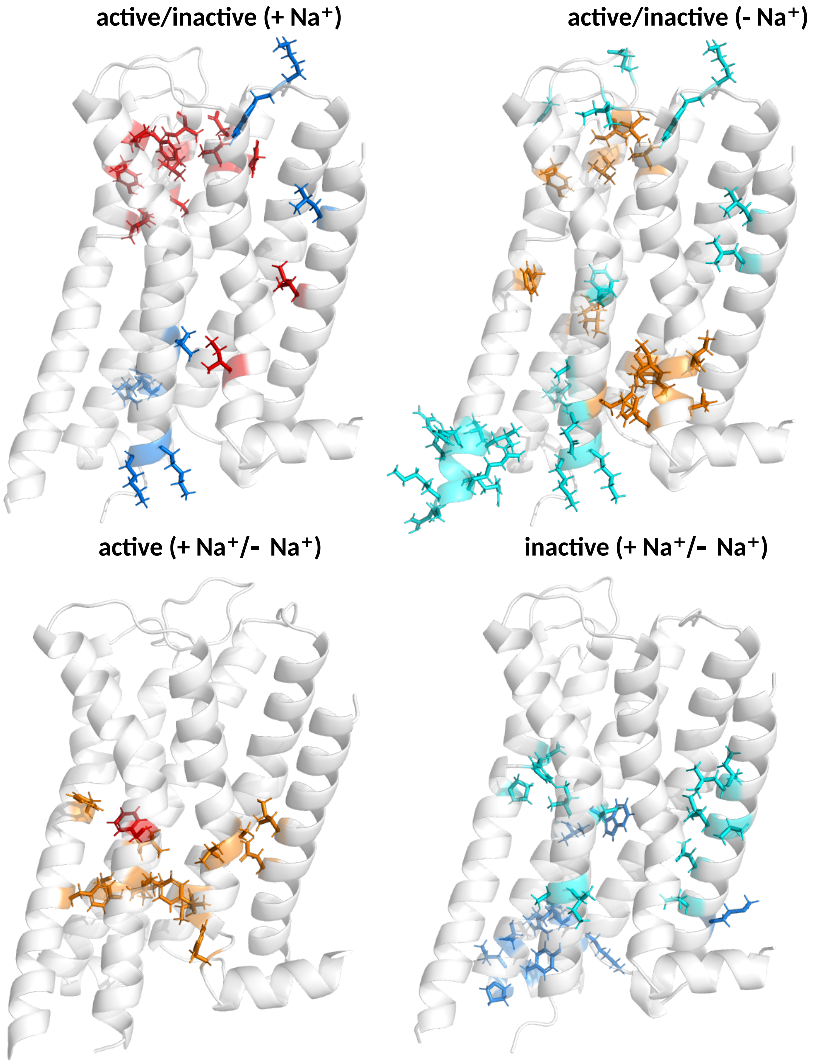


Figure 5: **Residue-wise water contact differences compared between two systems.** Water contacts enriched in active states are shown in red ($Na^+$-bound) and orange ($Na^+$-free); inactive-enriched contacts are shown in dark and light blue, respectively.

### References

(1) Selent, J.; Sanz, F.; Pastor, M.; De Fabritiis, G. Induced Effects of Sodium Ions on Dopaminergic G-Protein Coupled Receptors. *PLOS Computational Biology* **2010**, *6*, 1–6.

(2) Michino, M.; Free, R. B.; Doyle, T. B.; Sibley, D. R.; Shi, L. Structural basis for Na+-sensitivity in dopamine D2 and D3 receptors. *Chemical Communications* **2015**, *51*, 8618–8621.

(3) Chan, H. C. S.; Xu, Y.; Tan, L.; Vogel, H.; Cheng, J.; Wu, D.; Yuan, S. Enhancing the Signaling of GPCRs via Orthosteric Ions. *ACS Central Science* **2020**, *6*, 274–282.

(4) Neve, K. A.; Cumbay, M. G.; Thompson, K. R.; Yang, R.; Buck, D. C.; Watts, V. J.; DuRand, C. J.; Teeter, M. M. Modeling and Mutational Analysis of a Putative Sodium-Binding Pocket on the Dopamine D2 Receptor. *Molecular Pharmacology* **2001**, *60*, 373–381.

(5) Yang, D.; Zhou, Q.; Labroska, V.; Qin, S.; Darbalaei, S.; Wu, Y.; Yuliantie, E.; Xie, L.; Tao, H.; Cheng, J.; Liu, Q.; Zhao, S.; Shui, W.; Jiang, Y.; Wang, M.-W. G protein-coupled receptors: structure- and function-based drug discovery. *Signal Transduction and Targeted Therapy* **2021**, 2059–3635.
